# Whole Exome Sequencing identifies multiple pathogenic variants in a large south Indian family with Primary Open Angle Glaucoma

**DOI:** 10.1101/2020.09.21.306191

**Authors:** Mohd Hussain Shah, Manojkumar Kumaran, Prakash Chermakani, Mohideen Abdul Kader, R. Ramakrishnan, Subbiah. R. Krishnadas, Bharanidharan Devarajan, Periasamy Sundaresan

**Affiliations:** Department of Genetics, Aravind Medical Research Foundation, Madurai, Tamil Nadu, India; Department of Bioinformatics, Aravind Medical Research Foundation, Madurai, Tamil Nadu, India; Glaucoma Clinic, Aravind Eye Hospital, Tirunelveli, Tamil Nadu, India; Glaucoma Clinic, Aravind Eye Hospital, Madurai, Tamil Nadu, India; School of Chemical and Biotechnology, SASTRA (Deemed to be University), Thanjavur, Tamil Nadu, India; Department of Molecular Biology, Alagappa University, Karaikudi, Tamil Nadu, India

**Keywords:** Whole Exome Sequencing, Primary Open Angle Glaucoma, Genetic heterogeneity, South Indian, RPGRIP1 and ARGHEF40 gene

## Abstract

**Purpose:** To identify the pathogenic variants associated with POAG by using Whole Exome Sequencing (WES) data of a large South Indian family.

**Methods:** We recruited a large five generation of South Indian family (n=84) with positive family history of POAG. All study participants had comprehensive ocular evaluation (of the 84, 19 study subjects were diagnosed as POAG). Sanger sequencing of the candidate genes associated with POAG (*MYOC, OPTN and TBK1*) showed no genetic variation in the POAG affected family members. Therefore, we performed whole exome sequencing (WES) for 16 samples including (9 POAG and 7 unaffected controls) and the data was analysed using an in-house pipeline for prioritizing the pathogenic variants based on its segregation among the POAG individual.

**Results:** We identified one novel and five low-frequency pathogenic variants with consistent co-segregation in all affected individuals. The variant c.G3719A in RPGR-interacting domain of RPGRIP1 that segregated heterozygously with the six POAG cases is distinct from variants causing photoreceptor dystrophies, reported to affect the RPGR protein complex signaling in primary cilia. The cilia in TM cells has been reported to mediate the intraocular pressure (IOP) sensation. Furthermore, we identified a novel c.A1295G variant in Rho guanine nucleotide exchange factors Gene 40 (ARHGEF40) and likely pathogenic variant in the RPGR gene, suggesting that they may alter the RhoA activity essential for IOP regulation

**Conclusion:** Our study supports that low-frequency pathogenic variants in multiple genes and pathways probably affect the pathogenesis of Primary Open Angle Glaucoma in the large South Indian family.

## Introduction

Glaucoma is typically characterized by a progressive degeneration of optic nerve, which causes irreversible blindness. It is the second leading cause of global blindness after cataract ^[1]^. Primary open-angle glaucoma (POAG) is a subset of glaucoma majorly associated with loss of retinoganglion cells and their axons triggers permanent vision loss with an apprehensive exponential growth affecting around 60.5 million people worldwide. Due to the exponential increase in the global aging population, it is estimated that 80 million people will be affected by POAG by the end of 2020 ^[2]^ and the count could be expected to rise 111.8 million people by 2040. Which has an inexplicable impact on Asian and African peoples ^[3]^. Asia alone accounted for approximately 60% of the global glaucoma, whereas the Africa population represented (13%) is the second largest proportion of glaucoma cases in the world. To date in India, it is estimated that 12 million people have been affected by glaucoma and this number is expected to increase by 16 million by the end of 2020 ^[4, 5]^.

POAG is associated with several external risk factor including advanced age, central corneal thickness, myopia, steroid responsiveness and elevated intraocular pressure (IOP) ^[6]^. However, these risk factors do not capture the full spectrum of the disease. Though, the positive family history is also one of the risk factors for POAG. Genetic characterization of the POAG Positive family history, is useful for the identification of POAG-candidate genes (MYOC, OPTN, & TBK1) ^[6-8]^ that are capable of causing POAG. However, these candidate genes were discovered through large pedigrees with a positive family history of glaucoma. In addition, many studies have shown that POAG development is associated with various genetic risk factors, including genetic variants in CDKN2B-AS ^[9-12]^ CAV1/CAV2 ^[13]^, TMCO1 ^[14]^ AFAP1 ^[15]^, TXNRD2, FOXC1/GMDS, ATXN2 ^[16]^, FNDC3B ^[17, 18]^, GAS ^[14]^, PMM2 ^[19]^, TGFBR3 ^[20]^, and SIX1/SIX6 ^[10, 11]^. Due to its genetic heterogeneity and familial clustering nature of consistent genetic inheritance, necessitated an extensive molecular characterization to identify the factors responsible for the genetic predisposition in POAG affected individuals.

Our previous report, suggested that genetic screening of known candidate genes *(MYOC, OPTN* and *TBK1*) in a single large South Indian family with POAG did not detect the genetic risk factors underlying the pathogenesis of the disease ^[21]^. Therefore, this study aims to perform whole exome sequencing (WES), to identify the potential genetic risk factors associated with the positive POAG family history of the five-generation south Indian family.

## Methods

### POAG study subjects

The study was approved by the Institutional Review Board at the Aravind Eye Care System, Madurai, Tamil Nadu, India (IRB2011008BAS). This research adhered the tenets of the Declaration of Helsinki. All the study subjects were recruited and clinically evaluated as previously described ^[21]^. Briefly, an ophthalmic examination was conducted for 240 subjects during a field trip to Kayalpatanam, for this current study 84 members were recruited from a single family of a large South Indian family of five generations with a positive history of POAG.

### Whole exome sequencing

For Whole Exome Sequencing (WES) 5 ml of peripheral blood was collected from each study subjects. The genomic DNA was extracted using a salting-out precipitation method ^[22]^ and the concentration of the DNA samples was quantified using Qubit fluorometer. Samples were subjected for WES using the Agilent’s SureSelect Human All Exon V6 kit. The DNA libraries have been sequenced to mean >150X coverage on an Illumina HiSeq 4000 platform.

### Data Analysis

We developed an automated pipeline (Supplementary Figure.1) for the identification of pathogenic variant from WES data using UNIX script (https://github.com/bharani-lab/WES-pipelines/tree/master/Script). Raw reads (FASTQ file) were processed to remove the adapter and low-quality sequences using Cutadapt. Then the reads were further aligned against the human genome build GRCh37 using BWA-mem version 0.7.12. GATK version 4.1.0. for the identification of single-nucleotide variant (SNV) and small Insertion and Deletion (InDel) and it was further annotated using ANNOVAR ^[23]^. Variants were prioritized by setting the minor allele frequency (MAF) less than or equal to 0.5% in 1000genome, ESP, ExAC and genomeAD. For pathogenic variant identification, the non-synonymous, frameshift and splice variants were subsequently filtered by a two-step process; firstly, variants were prioritized based on their conservation score >2.5 (GERP score) and CADD score greater than 10; secondly, the variants should be predicted to be deleterious in at least three functional prediction tools (Polyphen2, SIFT, Mutation Taster, FATHMM and LRT). To avoid the mapping errors, all the variants were manually checked with IGV viewer. All predicted deleterious variants were further filtered based on their presence in at least more than three affected individuals in the pedigree. Finally, the variants were sorted out based on its segregation among their affected individuals. We used VarElect software ^[24]^ to sort the genes based on their direct or in direct association with glaucoma.

We performed pathway and gene ontology analysis using DAVID for all the genes identified in the final set of variants. A gene network was created using Cytoscape with the enriched pathways and biological processes.

### Sanger sequencing for the validation of novel variant

For segregation analysis, the novel variant of the ARHGEF40 gene was PCR amplified using the following gene specific primers (FW-5′-CTGAGCTGACGCCTGAACTT-3′); (RV-5′-GCCGTGGGTACTGAGAAAG-3′) and the fragments were bi-directional sequenced using (3130 Genetic Analyser; Applied Biosystems). Further the results were compared with the reference sequence of ARHGEF40 gene using NCBI-BLAST program and the chromatogram was analysed using Chromas lite (2.1) software.

## Results

### Clinical Evaluation of patients

A total of 84 family members were recruited from a single large south Indian family of five generation with a positive family history of POAG after a comprehensive ophthalmic screening of 240 family members in Kayalpatanam (as shown in the Supplementary Figure 2). Clinical assessment and complex pedigree analysis revealed that 19 of the 84 samples had been diagnosed with POAG. The clinical features of all 19 POAG affected individuals were discussed in detail ^[21]^. Only 9 out of 19 clinically diagnosed POAG individuals were subjected for Whole exome sequencing

### Exome Sequencing and Variant Filtering

Samples for the WES were selected solely on the basis of the POAG inheritance pattern observed in the pedigree of the South Indian family, which included nine POAG cases belonging to the generation (II-2,5,15; III-2,3,19,32 and IV-26,27) and seven unaffected controls (II-5; III-4,16,34,41 and IV-11,28) (Supplementary Figure 2). WES was carried out using Agilent SureSelect Human All Exon V6 kit and the DNA libraries have been sequenced to the Illumina HiSeq 4000 platform with an average coverage depth of ∼150x. The raw data were processed and analysed to identify the pathogenic variants (as indicated in the methods section of Figure 1). Approximately 60,000 variants (SNV and InDel) were identified in the exome of each patient’s aligned to the human reference genome build GRCh37.

**Figure 1.**
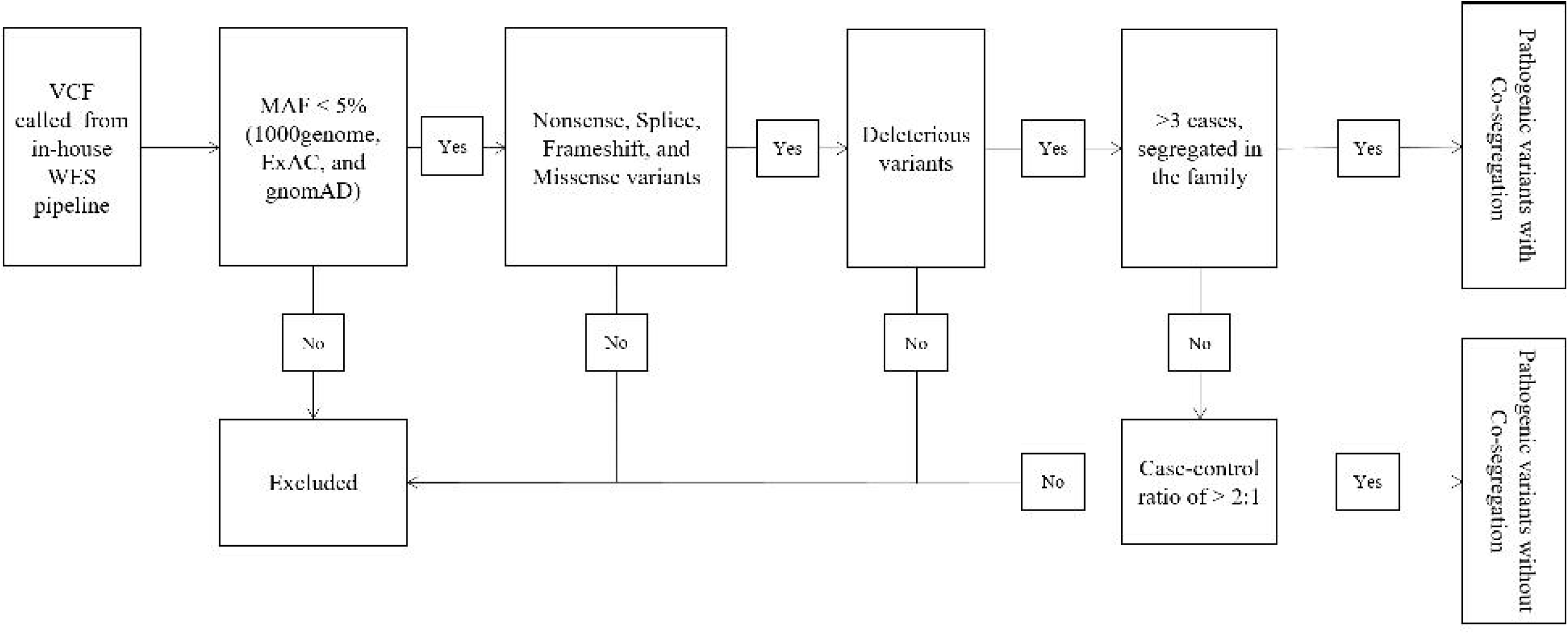
Work Flow for variant prioritization.

### Pathogenic variants

WES results revealed that pathogenic variants were identified in the six of the sixteen samples were co-segregated among the POAG affected family members (Supplementary Figure 2). Over all, we found six pathogenic variants (RPGRIP1; ARHGEF40; OR11H12; OR4K14; RNASE13 and OR11G2) (5 non-synonymous and one frameshift variant, respectively); all of which showed consistent co-segregation over generations in POAG affected individuals. Of the six pathogenic variants, RPGRIP1 alone showed direct association with glaucoma (Table.1). A heterozygous variant with MAF less than 0.01 in Retinitis Pigmentosa GTPase regulator-interacting protein1 (RPGRIP1) (c. G3719A; p. G1240E) had a deleterious effect exhibit highest phenotype score (Varelect score 8.35). Moreover, this variant also co-segregated in six POAG cases, which may affect the function of RPGR protein complex. In contrast, the RPGRIP1 variant (G1240E) is relatively common in most populations especially in Africans ^[25]^. A novel variant in Rho guanine nucleotide exchange factors Gene 40 ARHGEF40 gene (c. A1295G; p. Q432R), which was also segregated within the POAG affected individuals in the family. In addition, the novel variant of the ARHGEF40 gene was validated (c. A1295G; p. Q432R) in other family members through Sanger sequencing represented the genetic association in eight POAGs diagnosed and two unaffected family members (Supplementary Figure 2). Though the other four genetic variants of exome sequencing found consistent co-segregation OR11G2 (c.847delC; p. H282fs), OR4K14 (c. A355G; p. M119V), RNASE13 (c.C338T; p.S113F) and OR11H12 (c.T719G; p.V240G), but it did not show any association with glaucoma. In addition to these, this study also identified 54 pathogenic variants encoded by 51 different genes, but these genes are not co-segregated across the affected individuals. Among the 54 pathogenic variants, 52 were missense and 2 were Indel variants with frame-shifting coding regions as shown in Table 2. Based on the glaucoma clinical phenotype, the pathogenic variants, that are inconsistently segregating was also further prioritized in the POAG affected individual. The genetic variant in RPGR gene gene (c.G3430A; p.V1144I) was found in four POAG family members belongs to the generation (III-3;III-2;II-2;II-15) displayed it may affects its protein partner RPGRIP1 in the RPGR proteasome complex ^[26]^ and, eventually, PLK4 gene mutation in (c.C2513T; p.T838I) has adverse impact on microcephaly, failure of growth, and retinopathy ^[27]^. PLK4 mutations microcephaly, failure of growth, and retinopathy. Surprisingly, six of the 54 non-segregating pathogenic variants were identified as novel variants. Of which, the genetic variant in the neural cell adhesion molecule 1 (NCAM1) gene (c. A1841 T; p. D614V) was further confirmed by Sanger sequencing, since the genetic mutation in NCAM1 gene alters the optic nerve function which is further associated with increase in intraocular pressure^[28]^.

**Table 1.** List of the Pathogenic variants with co-segregation with phenotype. Varlect score with symbol † represents the direct association with glaucoma phenotypes and ‡ represent the indirect association. * represent the Novel variant.

| Chromosome<br>Position | Accession<br>number | Nucleotide<br>changes | Gene Name | Amino acid<br>change | dbSNP | Varlect | Number of cases (sample ID) |
| --- | --- | --- | --- | --- | --- | --- | --- |
| 14:21816432 | NM_020366.3 | c.G3719A | RPGRIP1 | p.G1240E | rs34725281 | 8.35† | 6 (III-3;III-2;II-2;III-32;IV-26;IV-27) |
| 14:21550588 | NM_001278529.2 | c.A1295G | ARHGEF40* | p.Q432R | . | 1.59‡ | 6 (III-3;III-2;II-2;III-32;IV-26;IV-27) |
| 14:20666340 | NM_001005503.1 | c.847delC | OR11G2 | p.H282fs | rs528205284 | 0.99‡ | 6 (III-3;III-2;II-2;III-32;IV-26;IV-27) |
| 14:20482998 | NM_001004712.1 | c.A355G | OR4K14 | p.M119V | rs7157076 | 0.95‡ | 6 (III-3;III-2;II-2;III-32;IV-26;IV-27) |
| 14:21502110 | NM_001012264.4 | c.C338T | RNASE13 | p.S113F | rs114504351 | 0.71‡ | 6 (III-3;III-2;II-2;III-32;IV-26;IV-27) |
| 14:19378312 | NM_001013354.1 | c.T719G | OR11H12 | p.V240G | rs61969158 | 0.22‡ | 6 (III-3;III-2;II-2;III-32;IV-26;IV-27) |

**Table 2.**
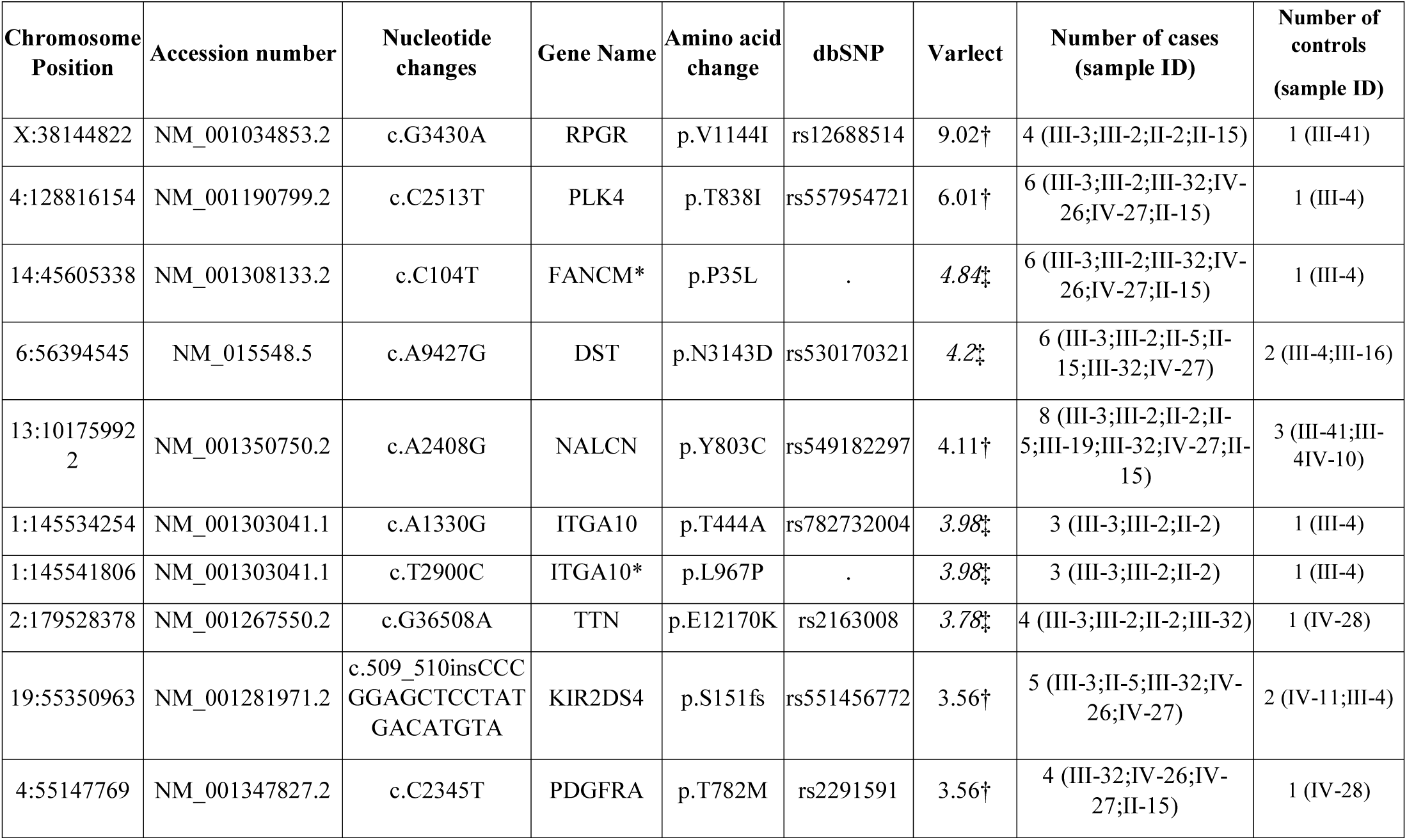
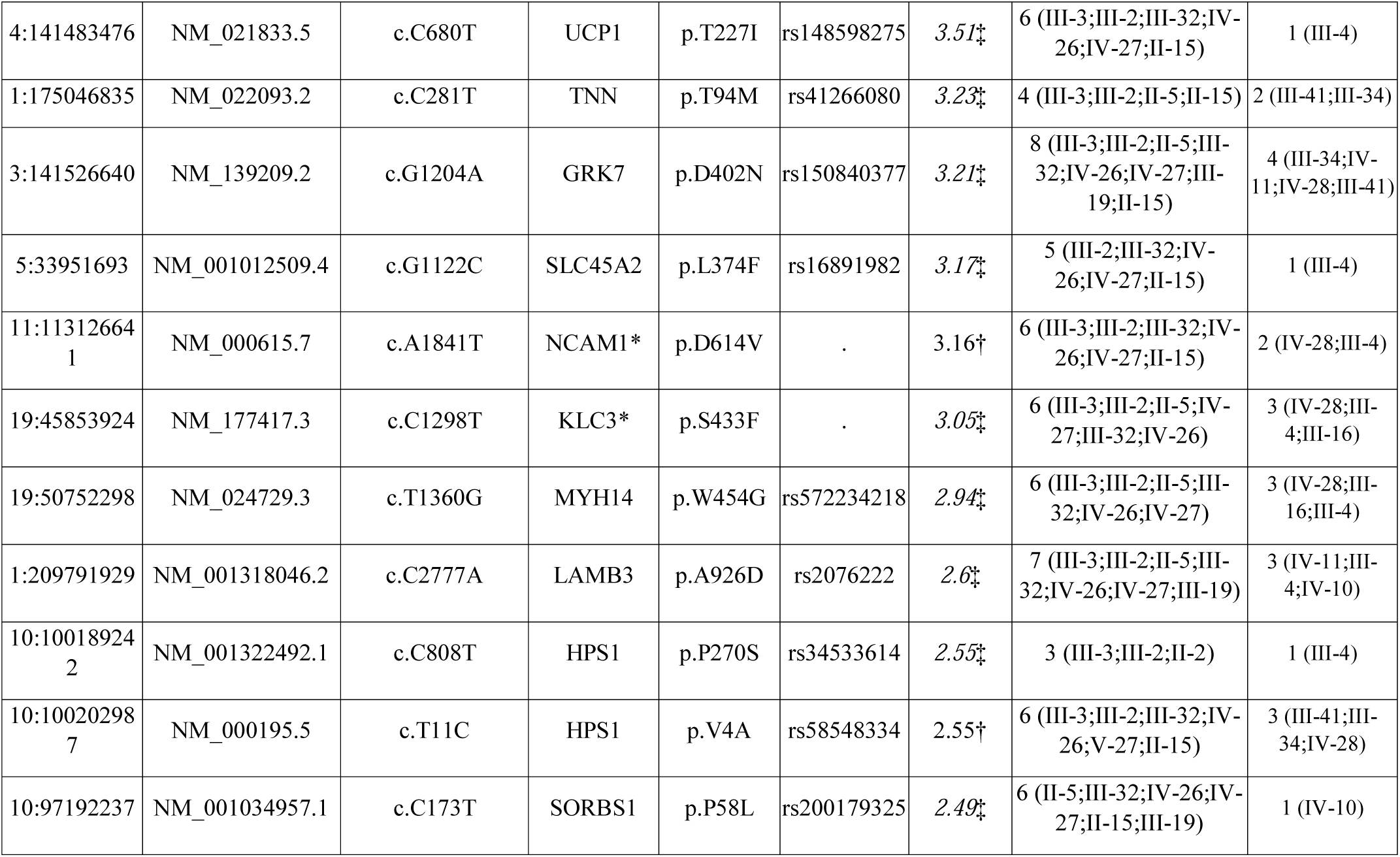
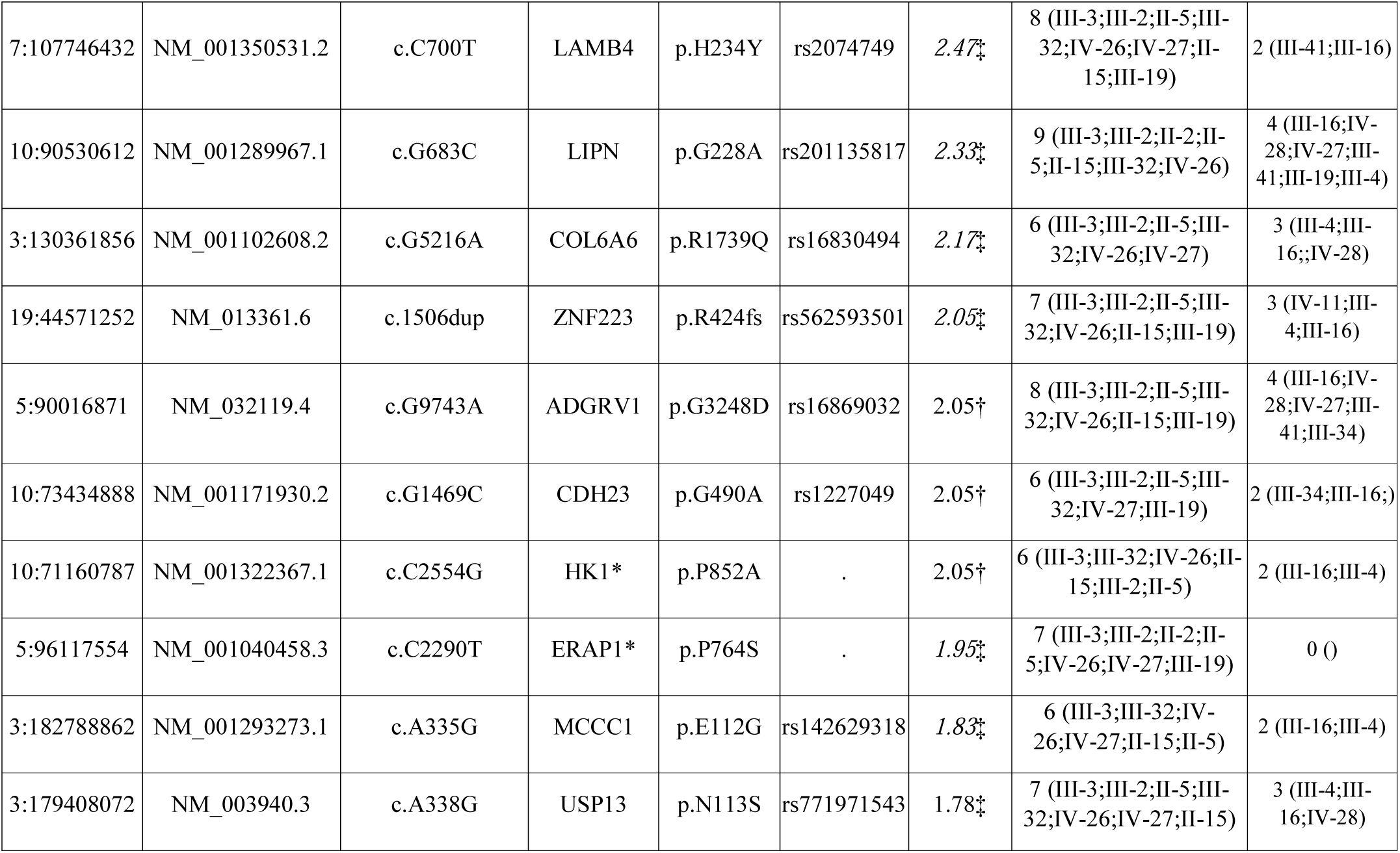
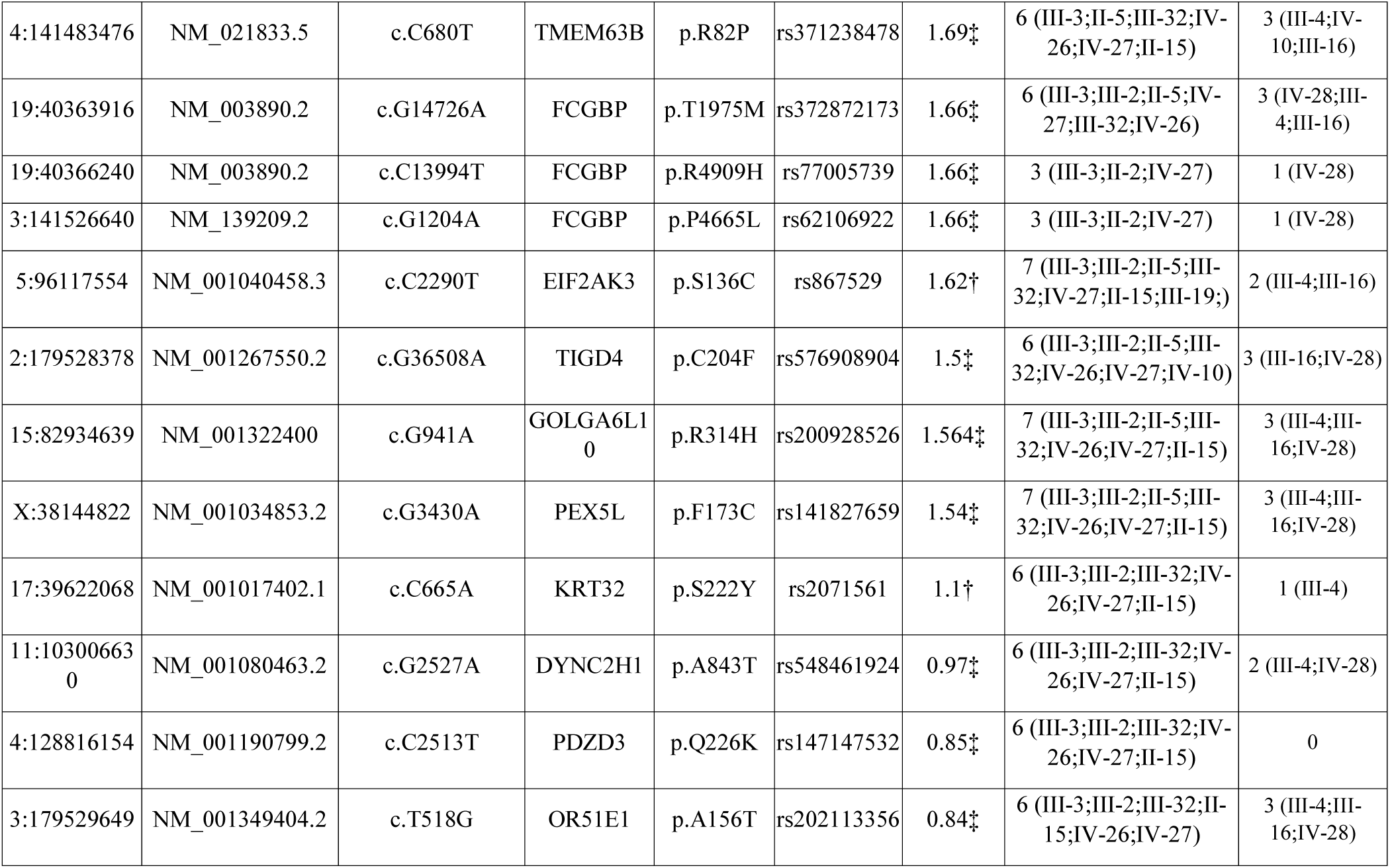
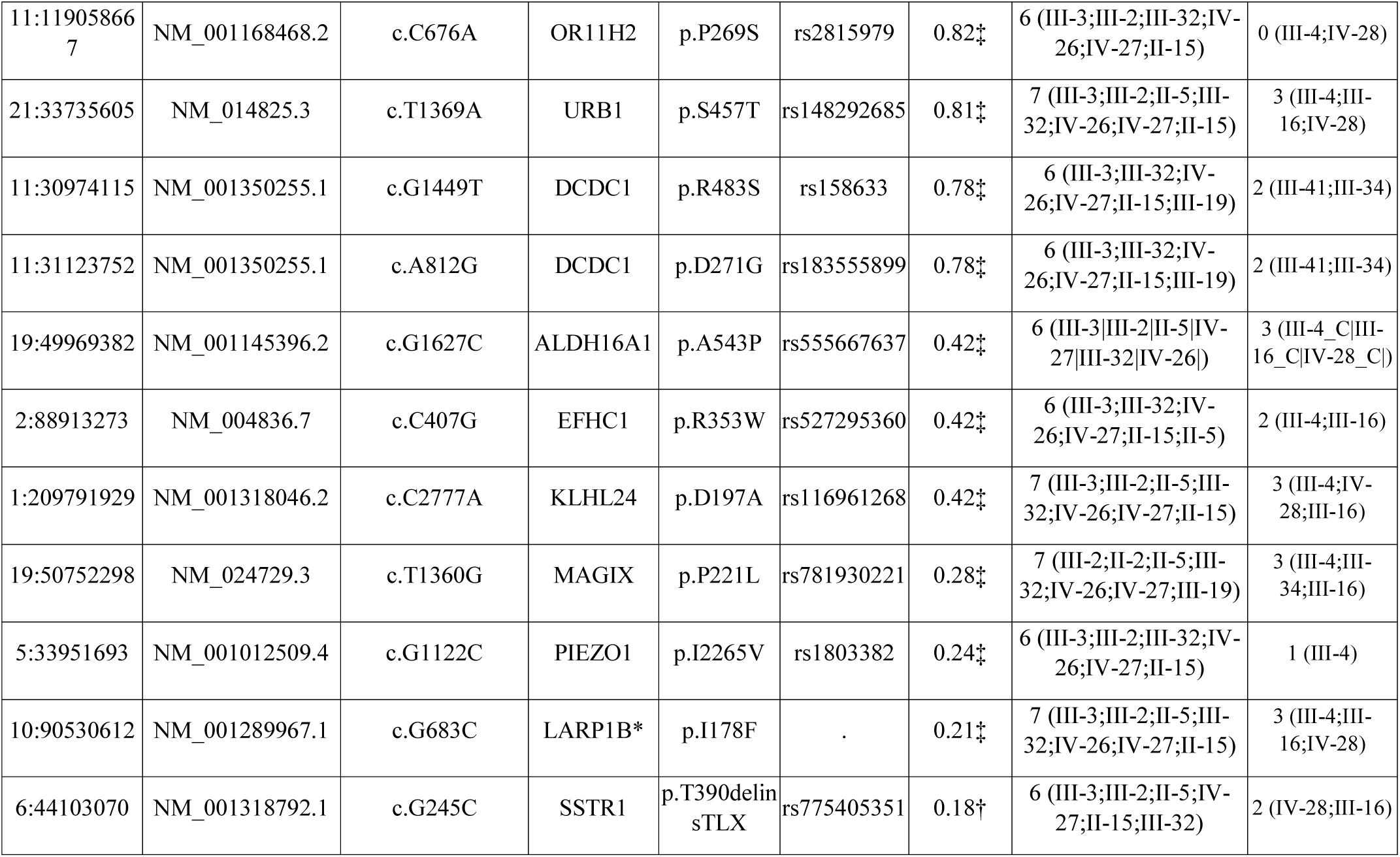
List of the Pathogenic variants without co-segregation with phenotype. Varlect score with symbol † represents the direct association with glaucoma phenotypes, and ‡ represent the indirect association. * represent the Novel variant.

### Functional network analysis

A functional network has been developed for all the pathogenic variants identified in POAG affected individuals, to investigate the pathways and biological processes involved in glaucoma pathogenesis. Initially, DAVID database was used to integrate all genes with KEGG pathways and Gene Ontology (GO) process. A total of 57 genes were significantly enhanced by three pathways, and 17 GO biological processes (P < 0.01). These pathways include Focal adhesion, ECM-receptor interaction and PI3K-Akt signalling pathway. Further Gene-functional network (Figure. 2), analysis revealed that NCAM1, LAMB4 and PDGFRA genes connected all the three pathways to the other GO process. Of these genes, NCAM1 was shown to be linked to RPGRIP1 and ARHGEF40 with pathogenic variants through RPGR protein interaction and GO processes of positive regulation of GTPase activity and visual perception.

**Figure 2.**
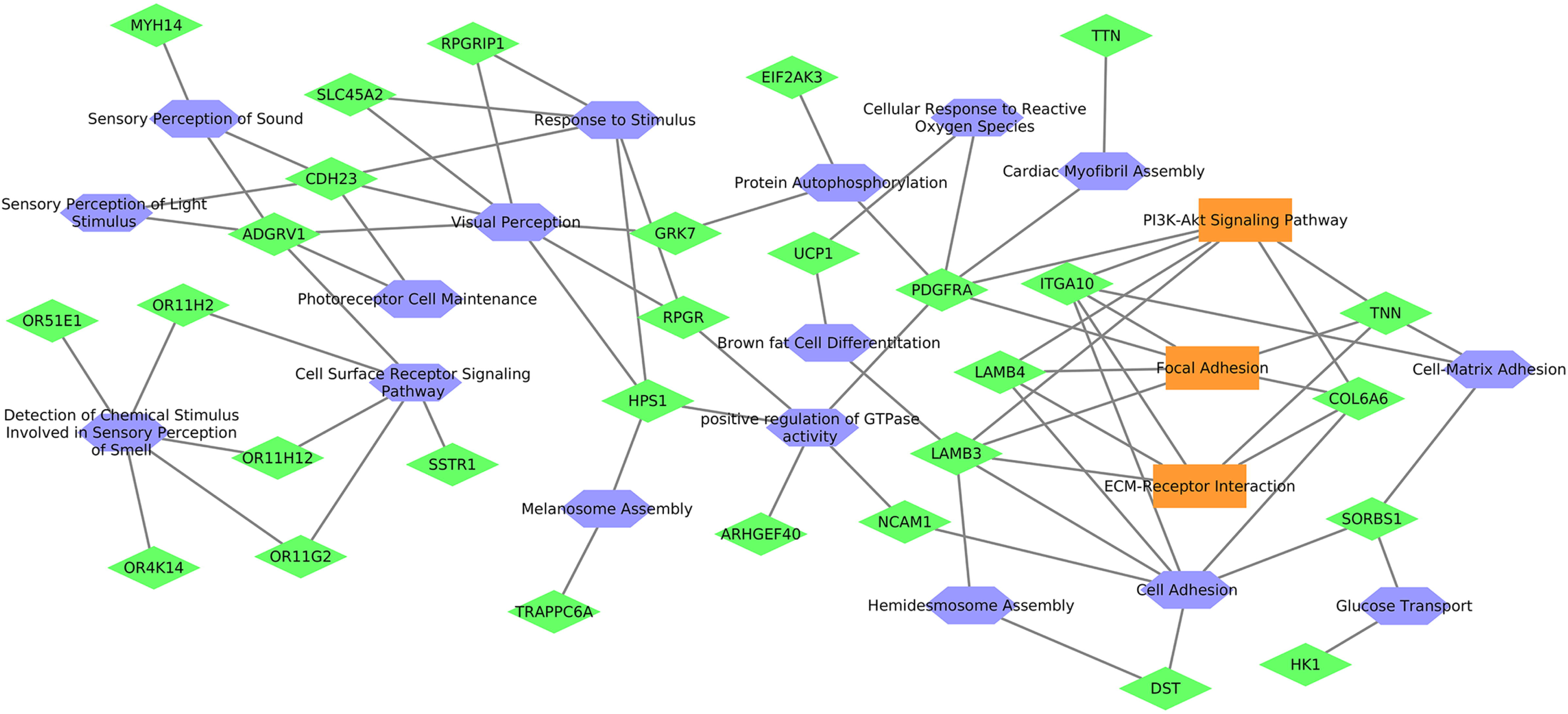
Functional network enriched with pathways and Gene Ontology (GO) on genes identified with pathogenic variants. Hexagon represents the gene; rectangle represent pathways and Diamond represents GO.

## Discussion

Studies of larger pedigree in POAG diagnosed families led to the discovery of mutation in MYOC, OPTN and TBK1genes. Ophthalmic examination of a single family in South India with 84 family members over five generations with Egyptian heritage had a positive POAG family history revealed no mutation for the primary candidate genes (MYOC, OPTN and TBK1) associated with POAG ^[21]^.

Hence WES of 16 samples including (9 POAG and 7 unaffected controls) of the 84 family members displayed a consistent co-segregation of six pathogenic genes ARHGEF40, RPGRIP1, OR4K14, RNASE13, OR11H12, and OR11G2 in six POAG samples. No pathogenic variants have been identified in 3 of the 9 POAGs and the remaining 7 unaffected individuals. Furthermore, candidate genes for the 3 individuals diagnosed with POAG can be identified through either deep intronic or whole genome sequencing. All the pathogenic variants identified from WES was further prioritized based on the glaucoma phenotype using an insilico VarElect phenotype sorting tool. Among the six co-segregating pathogenic variants, only two (ARHGEF40 and RPGRIP1) showed association with glaucoma. One of the key interesting fact is all six pathogenic variants were present in chromosome 14q, which had previously been reported to have potential POAG loci ^[10, 29, 30]^.

The pathogenic variant in retinitis pigmentosa GTPase regulator-interacting protein 1 (RPGRIP1) gene is observed with the highest phenotype score in six POAG cases, suggesting that it may have prominent role in POAG disease pathogenicity. Ferna’ndez-Marti’nez et al., has shown that the heterozygous non-synonymous variants in C2 domain of RPGRIP1gene might cause the various forms of glaucoma including POAG ^[25]^. In addition, it has also demonstrated that RPGRIP1 interaction with NPHP4 protein was shown to play a key role in glaucoma pathogenesis ^[25]^. In this study, four POAG cases were found to have a heterozygous pathogenic missense variant in the RPGR gene. In contrast to this study homozygous or compound heterozygous variants detected in RPGRIP1 are also associated to photoreceptor dystrophies ^[31, 32]^. Interestingly, we observed a pathogenic variant in RPGR gene, which is existed in four POAG cases. RPGRIP1 and its interacting partner RPGR, have been shown to express in human retina and also outside of the retina ^[26, 33-35]^ may regulate cilia genesis, maintenance, and function mainly through signalling pathways ^[36]^. Luo et al., 2014 has reported that the primary cilia of trabecular meshwork (TM) mediates intraocular pressure regulation through signalling pathway in the eye, and further highlighted that defect in the signalling pathway leads to Lowe syndrome that developed congenital glaucoma at birth ^[37]^. RPGR and its protein partners play an important role in actin cytoskeleton remodelling of cilia through these signalling pathways by activating the small GTPase, RhoA ^[38]^.

This research also identified a novel pathogenic variant in ARGHEF40 gene and this variant was further confirmed in all the affected family members using Sanger sequencing (Supplementary Figure 3). Study show that Rho guanine nucleotide exchange factors Gene Family protein (ARHGEF12) has been implicated as a risk factor of glaucoma by increasing intraocular pressure through RhoA/RhoA kinase pathway ^[39]^. Furthermore, the activation of the Rho/ROCK pathway results in trabecular meshwork (TM) contraction, and the inhibition of this pathway would aggravate relaxation of TM with a consequent increase in outflow facility and, thereby, decrease intraocular pressure ^[40]^. In the present study, we speculate that ARGHEF40 variant may affect the RhoA signalling through RPGRIP1 and its interacting partner RPGR in actin cytoskeleton remodelling of trabecular meshwork (TM) cilia, which may subsequently increase the intraocular pressure.

The pathogenic variants detected in other genes have not been reported to be directly associated with POAG. Therefore, we constructed a network of genes using GO and pathway enrichment. We have shown three pathways Focal adhesion, ECM-receptor interaction and PI3K-Akt signalling pathway to be associated with the pathogenesis of POAG. Furthermore, the highlighted genes ARHGEF40, RPGRIP1 and RPGR were enriched through visual perception and positive regulation of GTPase activity. Intriguingly, the genes NCAM1, HSP1 and PDGFRA including ARHGEF40 and RPGR in the biological process of positive regulation of GTPase activity is prioritized as top pathogenic variants based on the phenotype score. A study has shown that neural cell adhesion molecule (NCAM) participate in the optic nerve changes associated with elevated intraocular pressure ^[28]^.

## Conclusion

Overall, this study presented a panel of pathogenic variants in multiple genes, and their possible association with POAG pathogenesis in the five generation South Indian family. Thus, our findings strongly suggested that WES of the five-generation South Indian family showed extreme genetic heterogeneity of POAG within the family and the identified pathogenic variants showed continuous co-segregation among POAG affected individuals. Pathway analysis also displayed the association of the candidate genes involved in POAG pathogenesis but it requires larger sample size to ensure the authentic association of these identified genetic variants in POAG affected individuals.

## Acknowledgments

We thank all the patients and their families for their participation in this study. We are grateful to the Indian Council of Medical Research (ICMR) for their financial support. We are also thankful to Mr. Saravanan, Mrs. Kalarani and Mrs. Muthu Selvi for helping us in recruiting the samples and maintaining the clinical data in our lab.

## Ethics approval and consent to participate

The study adhered to the tenets of the Declaration of Helsinki, and ethics committee approval was obtained from the Institutional Review Board of the Aravind Eye Care System (IRB2011008BAS). All study participants read and signed informed consent after explaining the nature and possible significances of the study.

## Availability of data and material

The pipeline lines used for analysis in this study and a detailed tutorial is openly available for the public at (https://github.com/bharani-lab/Wole-Exome-Analysis-Pipeline).

We have submitted the data that support the finding of this study to SRA project ID PRJNA555016. Data can be accessed upon request

## Supportive/Supplementary Materials

### Supplementary Figure Legends

**Supplementary Figure 1.** Modular Pipeline

**Supplementary Figure 2.** Pedigree from south India Family. Family members diagnosed with POAG are shaded with black.

**Supplementary Figure 3.** Pedigree of selected family members from large south India family as shown in supplementary figure 1 (A). Sanger sequencing results of novel variant c.A1295G in ARGEF40 gene (marked with down arrow). The variant is detected in the family members II-2, III-1, III-2 and III-3 (B).

