## Supplementary figures and images for "Whole Exome Sequencing identifies multiple pathogenic variants in a large south Indian family with Primary Open Angle Glaucoma"

### Supplemental Figure 1

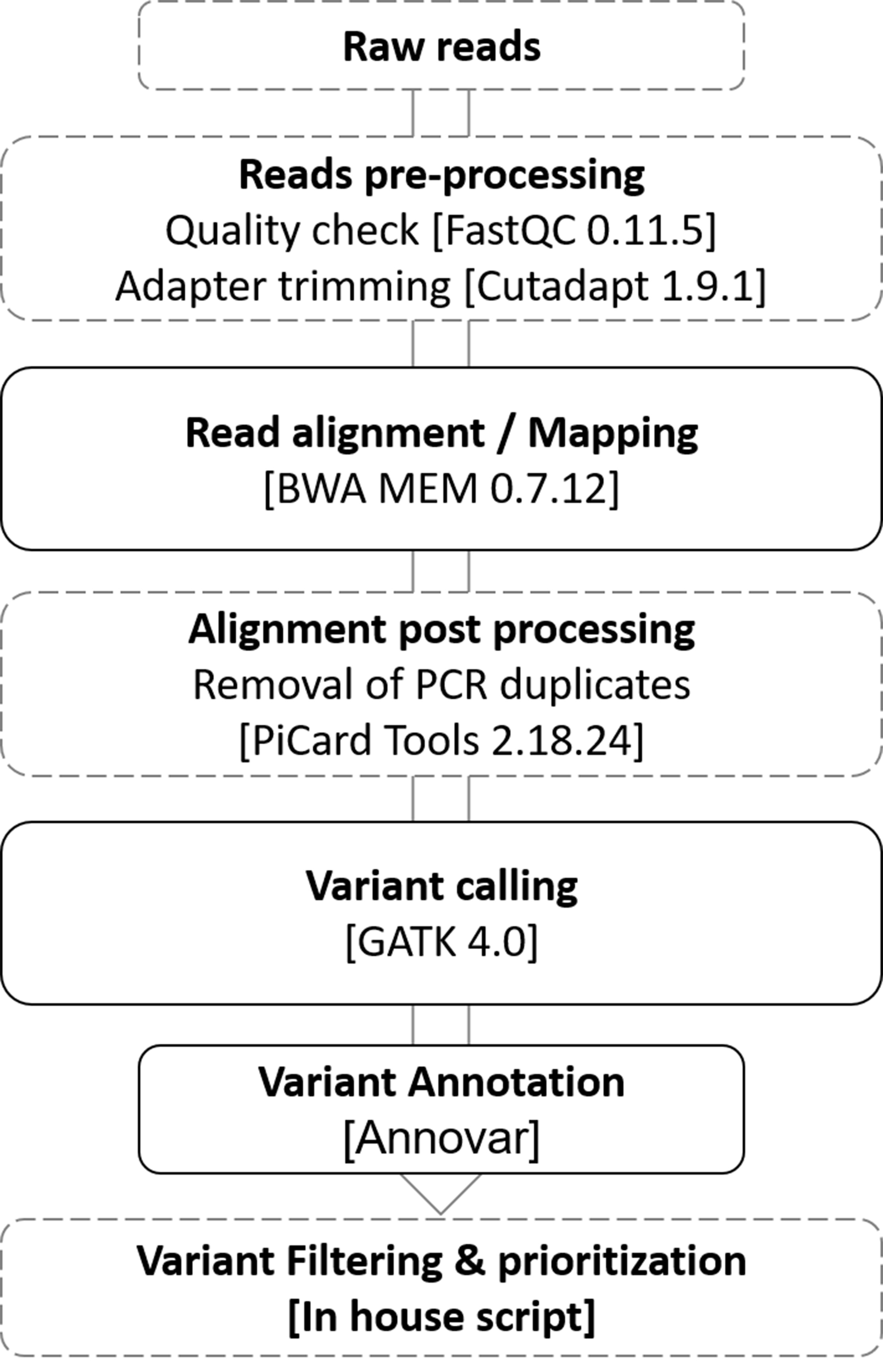

### Supplemental Figure 2

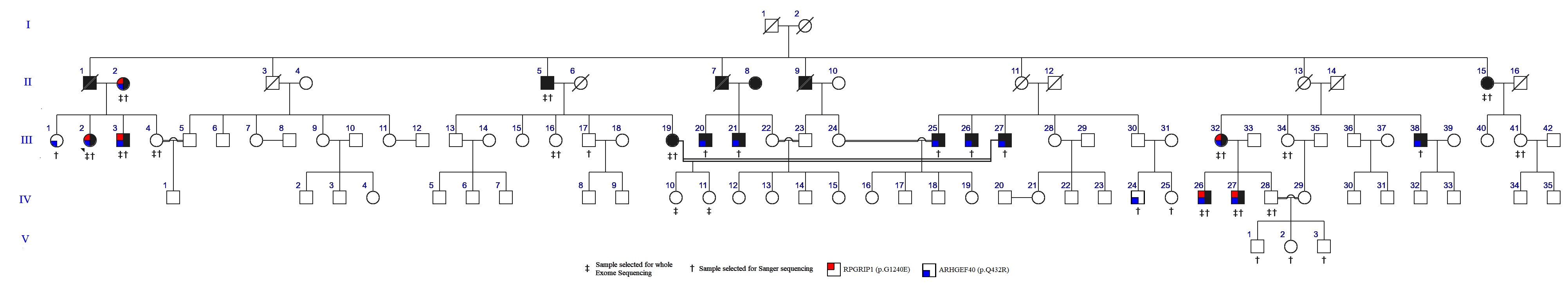

### Supplemental Figure 3

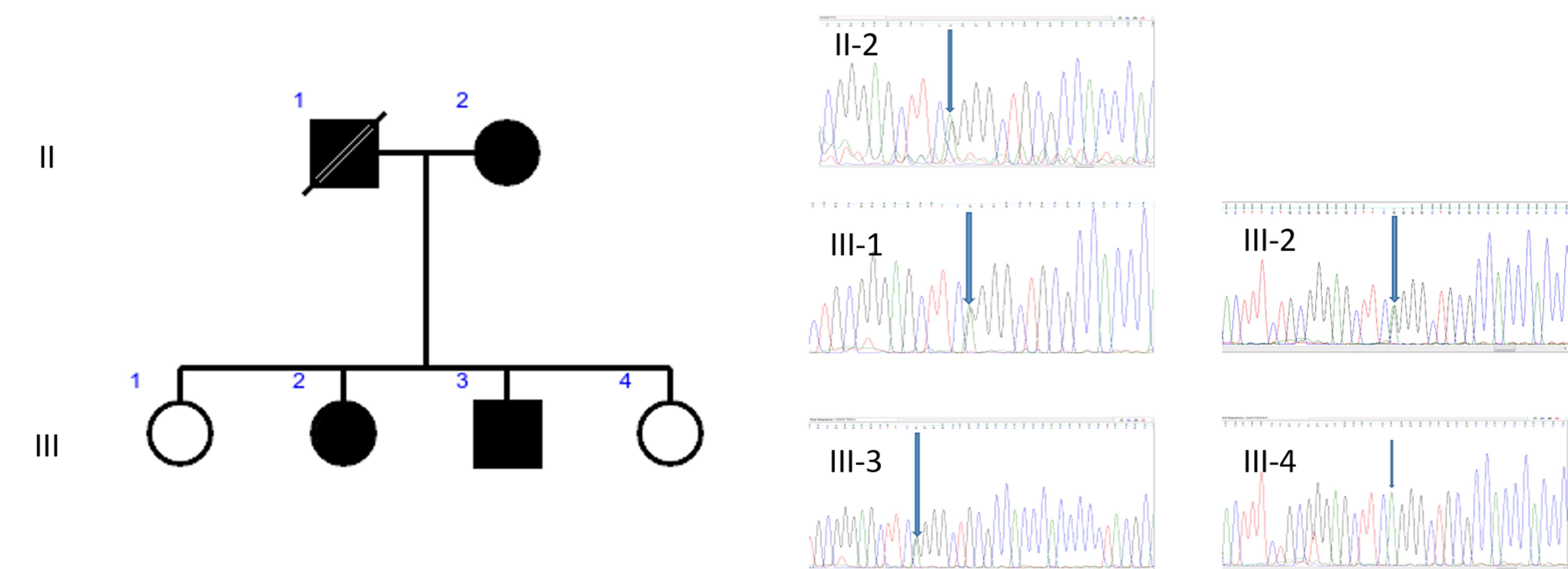
